# Accurate structure prediction of cyclic peptides containing unnatural amino acids using HighFold3

**DOI:** 10.1101/2025.03.06.641792

**Authors:** Sen Cao, Ning Zhu, Hongliang Duan

**Author notes:** To whom correspondence should be addressed: Hongliang Duan, Ning Zhu.

## Abstract

In recent years, cyclic peptides have emerged as a research hotspot in drug development due to their excellent stability, high specificity, and potential to penetrate intracellular targets. However, existing computational models still face challenges in making accurate predictions of the structures of cyclic peptides containing unnatural amino acids (unAAs), thereby limiting their application in drug design. The release of AlphaFold 3 (AF3) has significantly improved the modeling capability of biomolecular complexes and supports the definition of unAAs through residue modifications using the CCD database (Chemical Component Dictionary). Yet, its reliance on structures already present in the training library limits its ability to accurately predict cyclic peptide structures. Base on the AlphaFold 3 framework, we developed HighFold3, which comprises two sub-models: HighFold3-Linear and HighFold3-Cyclic, designed for predicting the structures of linear and cyclic peptides, respectively. Our research results show that, by incorporating a cyclic peptide positional preference matrix, HighFold3 achieves superior performance over other models (HighFold, HighFold2, and CyclicBoltz1) in predicting cyclic peptide structures, particularly excelling in complex cyclic peptides containing unAAs. HighFold3 provides an efficient tool for cyclic peptide drug design, promising to advance the application of cyclic peptides in targeted therapy, antibacterial treatments, and cellular penetration.

## 1. Introduction

Peptides have become a key focus in drug development due to their diverse functions and excellent pharmacological properties^1–3^. Cyclic peptides, as a unique class of peptides, exhibit higher stability, enhanced protein-protein interactions, and greater resistance to protease degradation than linear peptides due to their distinct cyclic structure^4–6^. This renders them highly attractive candidates for pharmaceutical research and drug development^7^. To improve their pharmacokinetic properties, enhance structural rigidity, and mitigate enzymatic degradation, cyclic peptides are often engineered to incorporate unAAs, thereby enhancing their potential as viable drug candidates^7,8^.

In recent years, protein structure prediction technologies have gained significant attention to accelerate the development of complex peptides, particularly cyclic peptides. The release of AlphaFold 2 revolutionized the modeling of protein structures and interactions^9^. With its new architecture and improved training methods, AlphaFold 3 further enhances the accuracy of predicting biomolecular complexes^10^. However, accurately predicting the structures of cyclic peptides unAAs remains a major challenge in computational biology^11^. Most deep learning-based structure prediction tools are trained primarily on existing structural data, which is predominantly derived from linear proteins^12^. Additionally, the same peptide sequence may adopt either a linear or cyclic conformation depending on environmental conditions or design goals, yet AlphaFold 3 remains limited in adapting to this conformational diversity^13^.

Meanwhile, novel tools for cyclic peptide structure prediction are continuously emerging. AFCyCDesign creatively integrates cyclic constraints into AlphaFold’s positional encoding, enabling the prediction of cyclic peptide structures^14^. Researchers combined high-T Molecular Dynamics (MD) with RSFF2C, accurately predicting 19 out of 23 cis-proline cyclic peptides with a backbone Root Mean Square Deviation (RMSD_Cα_) below 1 Å^15^. NCPepFold, based on the RFAA model, incorporates a cyclic positional matrix algorithm with the Transformer framework, effectively predicting cyclic peptide structures containing unAAs^13^. CyclicBoltz1, developed from Boltz-1 (an open-source variant of AlphaFold 3), specializes in predicting cyclic peptide models with all types of unAAs and demonstrates competitive performance against other tools. These evolving tools are unlocking new possibilities for cyclic peptide-based drug discovery^16,17^.

Building upon this foundation, we developed the HighFold series of models to enhance the accuracy and applicability of cyclic peptide structure prediction. HighFold modifies the positional matrices of AlphaFold and AlphaFold-Multimer, allowing for high-accuracy predictions of cyclic peptide monomers and their complexes, including those constrained by head-to-tail cyclization and disulfide bonds^18^. HighFold2, through fine-tuned training and the integration of atomic-level feature extraction, supports the structural modeling of cyclic peptides containing 23 commonly occurring unAAs, significantly improving the accuracy of spatial conformation modeling for cyclic peptides and their complexes with unAAs^11^. Expanding on these advancements, this study integrates cyclization constraints into AlphaFold 3, optimizing and introducing two sub-models tailored for different applications: HighFold3-Linear and HighFold3-Cyclic, designed for predicting the structures of linear and cyclic peptides containing unAAs, respectively. Additionally, HighFold3 utilizes chemical component data from the CCD to enhance unAA coverage^19^. Compared to existing cyclic peptide structure prediction tools, the HighFold3 series demonstrates superior accuracy and generalization capabilities, providing robust support for the development of complex peptide-based therapeutics.

## 2. Results and Discussion

We developed HighFold3 on the AlphaFold 3 framework to accurately predict the three-dimensional structures of peptide monomers and their complexes containing unAAs. The overall framework is shown in Figure 1A. HighFold3 introduces a cyclic positional matrix and employs a “Cyclization Switch” module to select the appropriate positional matrix based on the user-specified peptide type (linear or cyclic). Specifically, for linear peptides, the positional matrix defines the sequence interval between adjacent amino acids as 1 and the distance between the N-terminus and C-terminus as the peptide chain length minus 1 (Fig. 1B). For cyclic peptides, the positional matrix directly connects the N-terminus and C-terminus, forming a closed-loop structure (Fig. 1C). This design enables HighFold3 to predict distinct conformations for the same amino acid sequence based on user specifications. The selected positional matrix is encoded, combined with peptide feature data, and then fed into the AlphaFold 3 module for structure prediction. Additionally, HighFold3 is not limited by peptide chain length or conformational complexity, enabling efficient and high-precision structure predictions.

**Figure 1.**
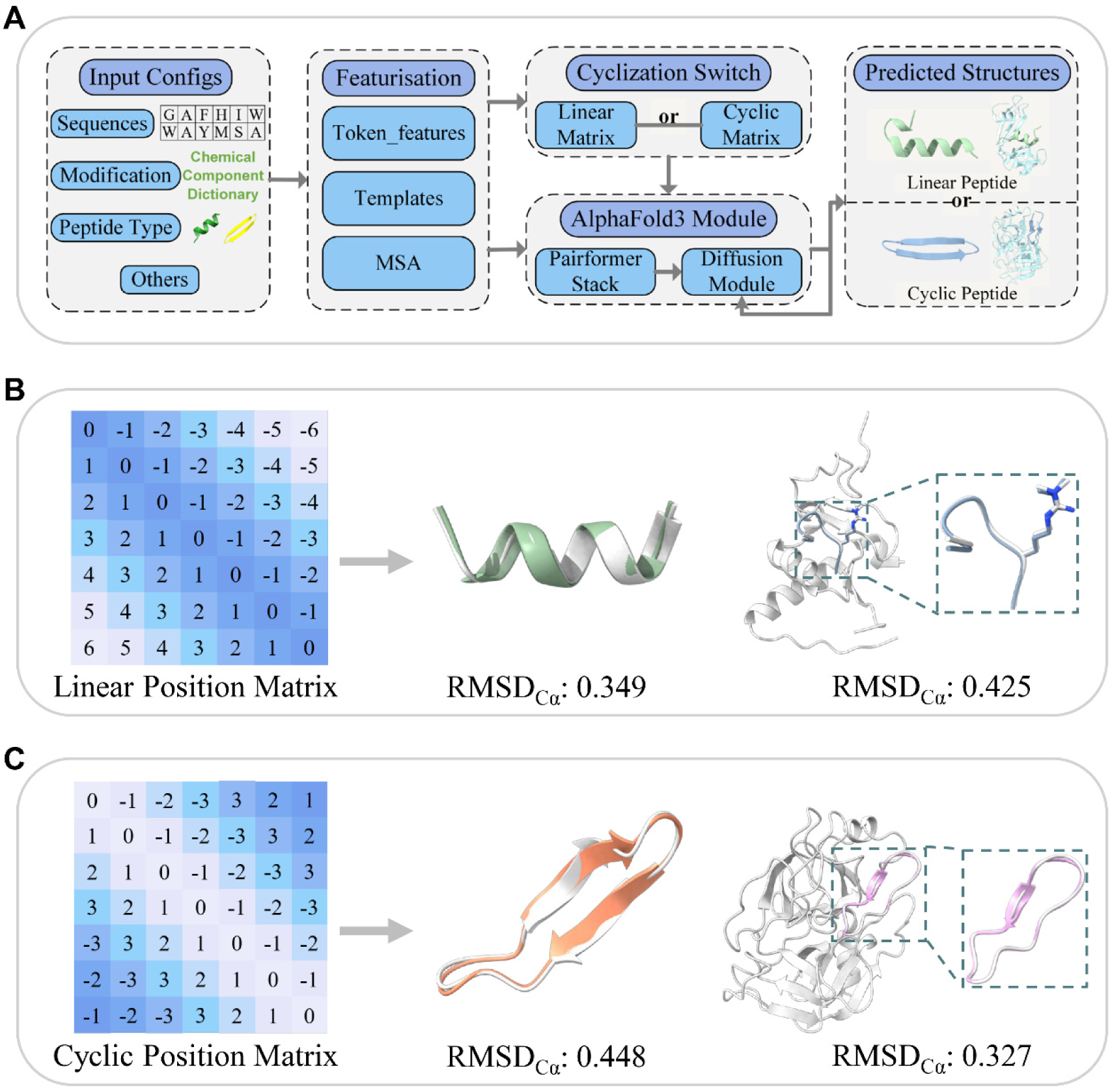
Overview of HighFold3. (A) A cyclization switch selects either a linear or cyclic position matrix based on user specifications. These features are processed through the AlphaFold 3 module, which consists of a Pairformer stack and a diffusion module, generating predicted structures for linear or cyclic peptides.(B) The linear positional matrix for linear peptides, where adjacent residues have a sequence interval of 1, and the N- and C-terminal residues are separated by a distance equal to the peptide length minus one. The predicted structure of a linear peptide is shown. (C) The cyclic positional matrix for cyclic peptides, where the N- and C-terminal residues are directly linked, forming a closed-loop structure. The predicted cyclic peptide structure is shown.

For performance evaluation, we employed RMSD as a key accuracy metric to systematically assess and compare HighFold3’s prediction capabilities. By default, we selected the Top-1 result from each prediction for evaluation. In the cyclic peptide test set, HighFold3 predicted 63 cyclic peptide samples from the AFCyCDesign dataset and compared the results with those of the HighFold and CyclicBoltz1 models. Additionally, HighFold3 predicted 18 cyclic peptide samples containing unAAs and cyclized via head-to-tail amide bonds from the HighFold2 dataset, with results compared to the HighFold2 and CyclicBoltz1 models. In the linear peptide test set, the HighFold3-Linear module predicted 36 linear peptide samples containing unAAs from the HighFold2 dataset, with results compared to the HighFold2 model.

To further evaluate the model’s versatility, we selected a diverse set of sample sequences from the CyclicPepedia cyclic peptide database, the CPPsite linear peptide database, and the ParaPep linear peptide database^20–22^. These samples encompass a variety of sequence lengths and amino acid compositions (see the “Materials and Methods” section for details).These samples included 100 cyclic peptide sequences, 50 cyclic peptide sequences with unAAs, 100 linear peptide sequences, and 50 linear peptide sequences with unAAs. To comprehensively evaluate the accuracy and robustness of HighFold3-Linear and HighFold3-Cyclic across different peptide types, we introduced “cyclization rate” and “non-cyclization rate” as additional evaluation metrics.

### 2.1 Accurate structure prediction of Cyclic peptides

We evaluated 63 cyclic peptide monomer structures from the AFCyCDesign dataset, using RMSD_Cα_ as a key metric to quantify the deviation between predicted and experimental structures. We benchmarked HighFold3’s predictions against HighFold and CyclicBoltz1 models to comprehensively assess its predictive performance.

As shown in Figure 2A, HighFold3 demonstrates superior prediction accuracy: its median RMSD_Cα_ is 0.918 Å, and the average RMSD_Cα_ is 1.337 Å. In comparison, HighFold has a median RMSD_Cα_ of 1.058 Å and an average RMSD_Cα_ of 1.478 Å, reflecting improvements of 13.23% and 9.54%, respectively. Among the 63 test samples, HighFold3 outperforms HighFold in 36 cases, whereas CyclicBoltz1 surpasses HighFold in only 32 samples. Notably, when compared to natural structures, HighFold3 achieves RMSD_Cα_ below 1.5 Å in 44 out of 63 samples, significantly exceeding HighFold (37 samples) and CyclicBoltz1 (38 samples) under the same metric. These findings indicate that HighFold3 achieves near-experimental accuracy in predicting cyclic peptide monomer structures, closely approaching high-resolution X-ray crystallography and NMR results.

**Figure 2.**
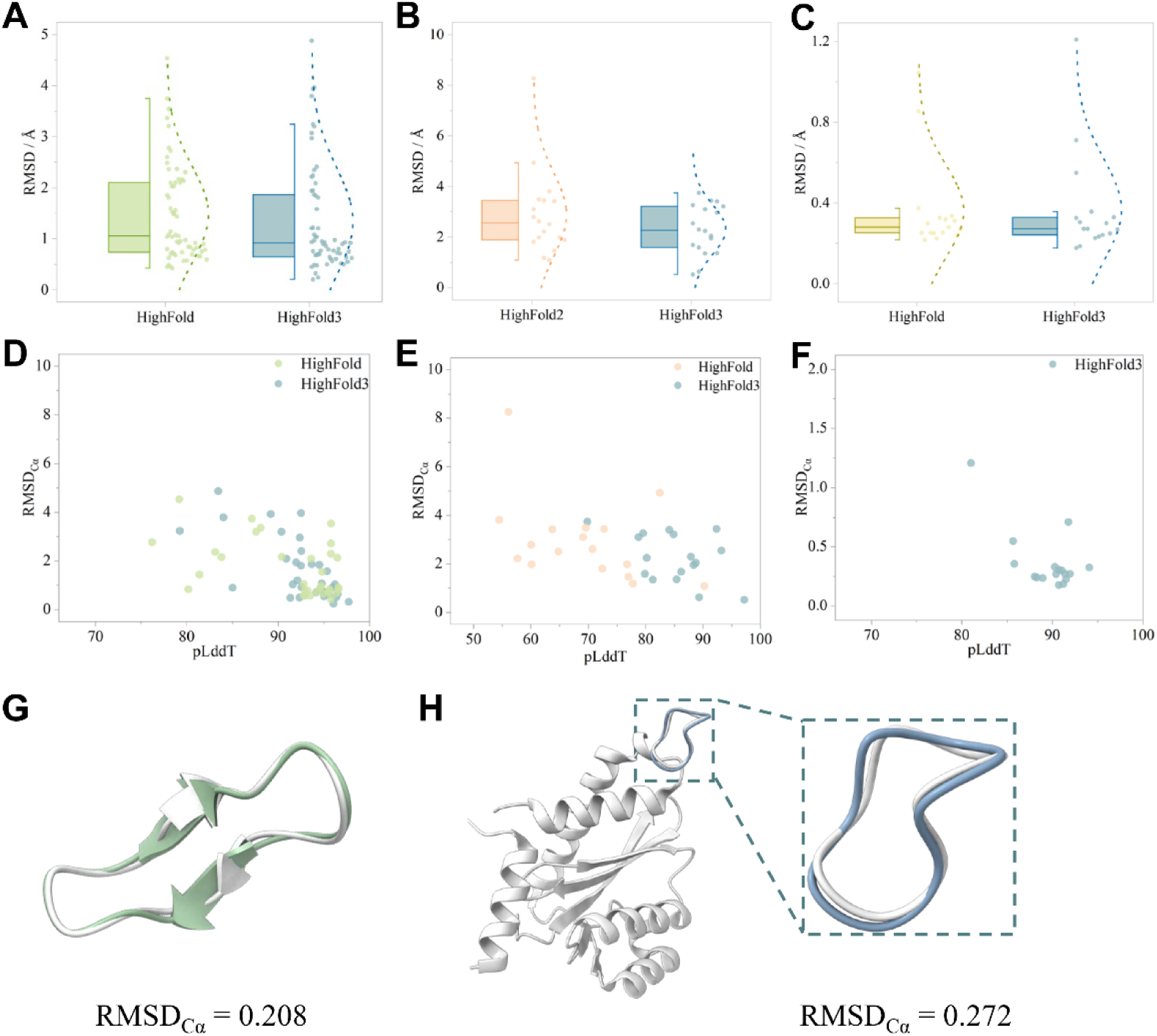
Performance Evaluation of HighFold3 in Predicting Cyclic Peptide Monomer and Complex Structures. (A) RMSD_Cα_ distribution of cyclic peptide samples predicted by HighFold and HighFold3. (B) RMSD_Cα_ distribution of cyclic peptide samples containing unAAs predicted by HighFold2 and HighFold3. (C) RMSD_Cα_ distribution of cyclic peptide complex samples predicted by HighFold and HighFold3. (D) Correlation between peptide pLDDT and RMSD_Cα_ in cyclic peptide samples predicted by HighFold and HighFold3. (E) Correlation between peptide pLDDT and RMSD_Cα_ in cyclic peptide samples containing unAAs predicted by HighFold2 and HighFold3. (F) Correlation between peptide pLDDT and RMSD_Cα_ in cyclic peptide complex samples predicted by HighFold3. (G) Example of cyclic peptide monomer prediction. The native structure is shown in gray, and the predicted structure in green. (H) Example of cyclic peptide complex prediction. The native structure is shown in gray, and the predicted structure in blue.

To further evaluate HighFold3’s ability to predict complex structures, we tested 17 head-to-tail cyclized peptide monomers containing unAAs from the HighFold2 dataset. We employed RMSD_Cα_, all-atom RMSD (RMSD_all-atom_), and RMSD of unnatural amino acids (RMSD_unAA_) as key metrics. These metrics comprehensively capture the model’s accuracy in predicting complex cyclic peptide structures, particularly in modeling unAA side-chain conformations. As shown in Figure 2B, HighFold3 demonstrates strong performance: its median RMSD_Cα_ is 2.299 Å, and the average RMSD_Cα_ is 2.342 Å. In comparison, HighFold2 has a median RMSD_Cα_ of 2.791 Å and an average RMSD_Cα_ of 2.909 Å, reflecting improvements of 17.63% and 19.49%, respectively. Among these 17 test cases, HighFold3 outperforms HighFold2 in 10 instances, demonstrating its robustness in modeling complex cyclic peptides and unAAs.

In practical cyclic peptide binder design, modeling the monomer alone is often insufficient. To develop potential binders, we need to predict the full complex structure, including the target protein and its cyclic peptide ligand. We evaluated 17 complex structures from the HighFold dataset, where cyclic peptide ligands were defined using the cyclic position matrix and assessed using ligand RMSD_Cα_. As shown in Figure 2C, HighFold3 outperforms HighFold in 10 out of 17 test samples.These findings indicate that HighFold3 not only achieves high accuracy in monomer prediction but also effectively models interactions between cyclic peptides and proteins, providing robust support for complex modeling in drug design.

To comprehensively assess the reliability of HighFold3’s predictions, we examined the correlation between confidence scores (pLDDT) and RMSD_Cα_ across three test sets. In the 63 cyclic peptide monomers from the AFCyCDesign dataset, RMSD_Cα_ exhibits a negative correlation with pLDDT, with a Pearson correlation coefficient of -0.67. This indicates that higher pLDDT values generally correspond to lower RMSD, confirming a direct relationship between model confidence and prediction accuracy. Notably, among HighFold3’s high-confidence predictions (pLDDT > 0.85), 44 samples achieve RMSD_Cα_ below 1.5 Å, further demonstrating its crystal-like precision. In the unAA-containing cyclic peptide monomer test set, RMSD_Cα_ also shows a negative correlation with pLDDT, with a Pearson coefficient of -0.45. Among HighFold3’s 17 samples, 6 exhibit RMSD < 2 Å at high confidence (pLDDT > 0.70), compared to 5 for HighFold, indicating HighFold3’s superior adaptability in unAA modeling. In the cyclic peptide complex test set, the correlation coefficient is -0.66, further confirming the model’s stability and consistency. These findings demonstrate that HighFold3 makes significant strides in predicting complex cyclic peptide structures, including those with unAAs.

Finally, we visualized HighFold3’s outstanding capability in modeling cyclic peptide monomers and complex structures. As shown in Figure 2G, for the PawS-derived peptide (PDB ID: 2LWU) monomer structure, HighFold3 achieves high accuracy with an RMSD_Cα_ of just 0.208 Å, indicating that the model can precisely reproduce the atomic-level conformation of natural cyclic peptides. Additionally, HighFold3 excels in predicting the complex structure of HIV integrase with an allosteric peptide inhibitor (PDB ID: 3AVB), where the ligand cyclic peptide’s RMSD_Cα_ is 0.272 Å (Figure 2H). This accuracy validates HighFold3’s excellent modeling capabilities in cyclic peptide complex prediction.

The above results indicate that HighFold3 demonstrates significant advantages in predicting cyclic peptide monomers and complexes containing unAAs, providing a high-precision modeling tool for peptide drug design.

### 2.2 Accurate structure prediction of Linear peptides

We extended our evaluation of HighFold3’s predictive performance to a linear peptide test set containing unAAs, assessing both monomer structures and their complexes with proteins. The evaluation metrics included RMSD_Cα_, RMSD_all-atom_, and RMSD_unAA_ to comprehensively assess the model’s accuracy in predicting backbone and side-chain conformations.

As shown in Figure 3A, in the linear peptide test set, HighFold3 achieves an average RMSD_Cα_ of 1.342 Å and a median of 0.979 Å, whereas HighFold2 has an average RMSD_Cα_ of 2.080 Å and a median of 0.994 Å, demonstrating improvements of 35.48% and 1.51%, respectively. Figure 3B indicates that HighFold3’s average RMSD_all-atom_ is 1.710 Å with a median of 1.373 Å, compared to HighFold2’s average of 3.037 Å and median of 1.906 Å, reflecting improvements of 43.7% and 28%, respectively. Additionally, Figure 3C reveals that HighFold3’s average RMSD_unAA_ is 1.021 Å with a median of 1.042 Å, whereas HighFold2’s average is 2.826 Å and median is 1.971 Å, showing significant enhancements of 63.9% and 47.1%, respectively. These findings strongly validate HighFold3’s superior capability in modeling the three-dimensional structures of linear peptide monomers and complexes containing unAAs, particularly excelling in capturing unAA side-chain conformations with high precision. To comprehensively evaluate the model’s prediction reliability, we examined the correlation between the three evaluation metrics (RMSD_Cα_, RMSD_all-atom_, RMSD_unAA_) and pLDDT in the test set. As shown in Figure 3D, pLDDT for linear peptides exhibits a strong negative correlation with RMSD_Cα_, with a Pearson correlation coefficient of -0.58. In HighFold3’s high-confidence predictions (pLDDT > 0.85), 24 out of 36 samples have an RMSD_Cα_ below 1.5 Å, indicating that the model can reliably predict backbone structures at high confidence. Figure 3E reveals that the Pearson correlation coefficient between pLDDT and RMSD_all-atom_ is -0.59, and at the same high confidence level (pLDDT > 0.85), 19 out of 36 samples achieve an RMSD_all-atom_ below 1.5 Å. This highlights not only the accuracy of backbone prediction but also HighFold3’s exceptional precision in modeling side-chain conformations. Additionally, as shown in Figure 3F, the Pearson correlation coefficient between pLDDT and RMSD_unAA_ for unAAs is -0.51. Among HighFold3’s 36 samples, 27 exhibit an RMSD_unAA_ below 2 Å at good confidence (pLDDT > 0.70), compared to only 16 for HighFold2, underscoring HighFold3’s enhanced adaptability in unAA structural modeling.

**Figure 3.**
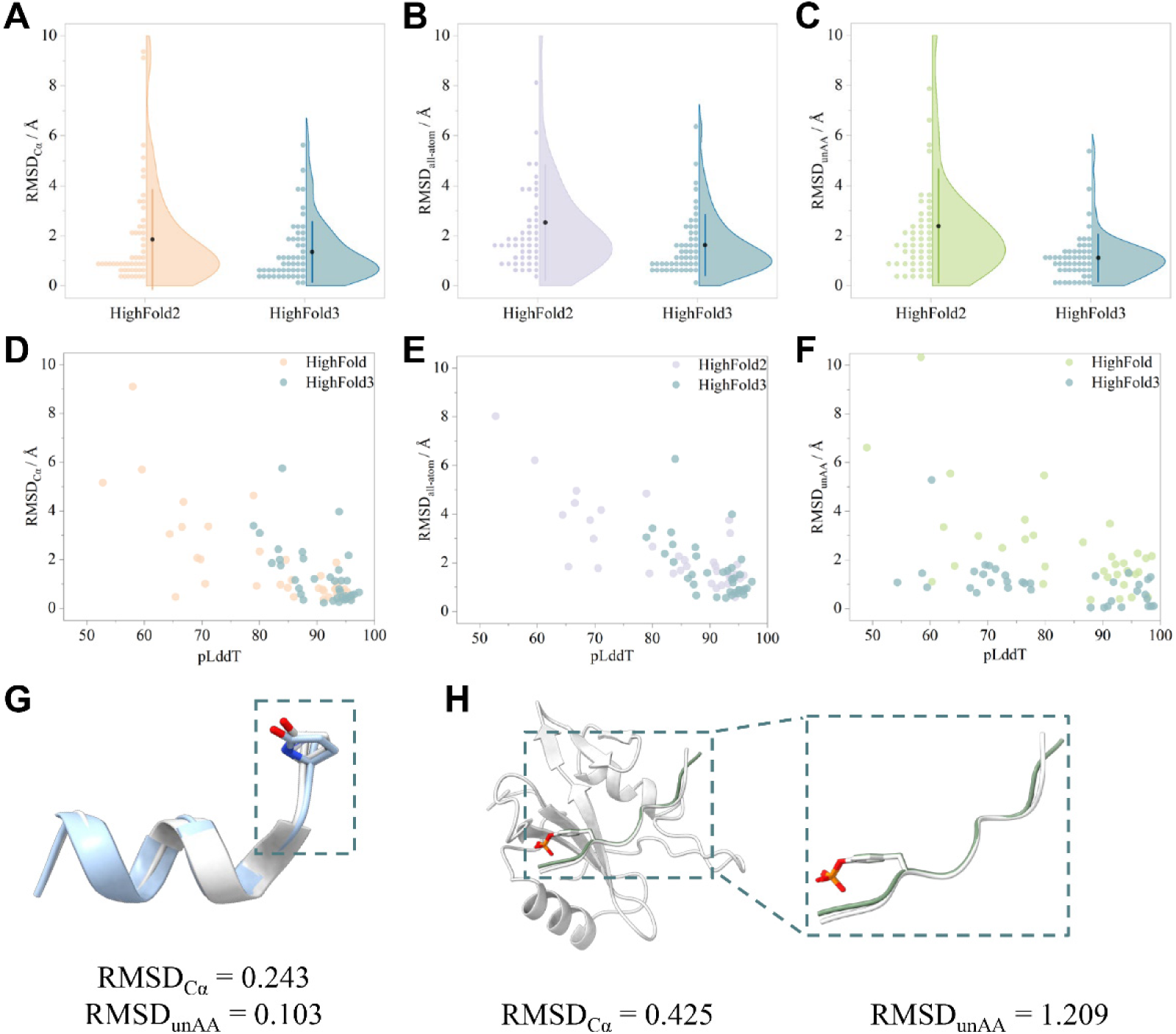
Performance Evaluation of HighFold3 in Predicting Linear Peptide Monomer and Complex Structures (A) RMSD_Cα_ distribution of linear peptide samples predicted by HighFold2 and HighFold3. (B) RMSD_all-atom_ distribution of linear peptide samples predicted by HighFold2 and HighFold3. (C) RMSD_unAA_ distribution of linear peptide samples predicted by HighFold2 and HighFold3. (D) Correlation between peptide pLDDT and RMSD_Cα_ in linear peptide samples predicted by HighFold2 and HighFold3. (E) Correlation between peptide pLDDT and RMSD_all-atom_ in linear peptide samples predicted by HighFold2 and HighFold3. (F) Correlation between peptide pLDDT and RMSD_unAA_ in linear peptide samples predicted by HighFold2 and HighFold3. (G) Example of linear peptide monomer prediction. The native structure is shown in gray, and the predicted structure in light blue. (H) Example of linear peptide complex prediction. The native structure is shown in gray, and the predicted structure in cyan.

### 2.3 Advantages of HighFold3 in Structure Prediction

While HighFold3 demonstrates significant improvements in cyclic peptide structure prediction, part of this progress may stem from the extensive cyclic peptide structures in the training data of its underlying framework, AlphaFold 3. However, we observed that AlphaFold 3 struggles with predicting cyclic peptide structures, particularly for unseen cyclic peptide sequences. AlphaFold 3 lacks specific constraints for closed-loop structures, which can lead to poor prediction performance. To address this issue, we introduced a cyclic peptide position matrix into the AlphaFold 3 framework to enhance its ability to model cyclic peptide conformations. This modification aimed to evaluate whether HighFold3 can overcome this limitation and enhance generalization.

To test this hypothesis, we selected a head-to-tail cyclic peptide structure (PDB ID: 9HVC) that was absent from AlphaFold 3’s training set and predicted it using both AlphaFold 3 and HighFold3^23^. As shown in Figure 4A and Figure 4B, AlphaFold 3 generates a linear peptide conformation for this cyclic sequence, with an N-to-C terminal distance exceeding 20 Å, whereas HighFold3 accurately predicts a closed-loop structure. This result highlights AlphaFold 3’s lack of specific constraints for cyclic structures, while HighFold3 enhances the model’s generalization capability.

**Figure 4.**
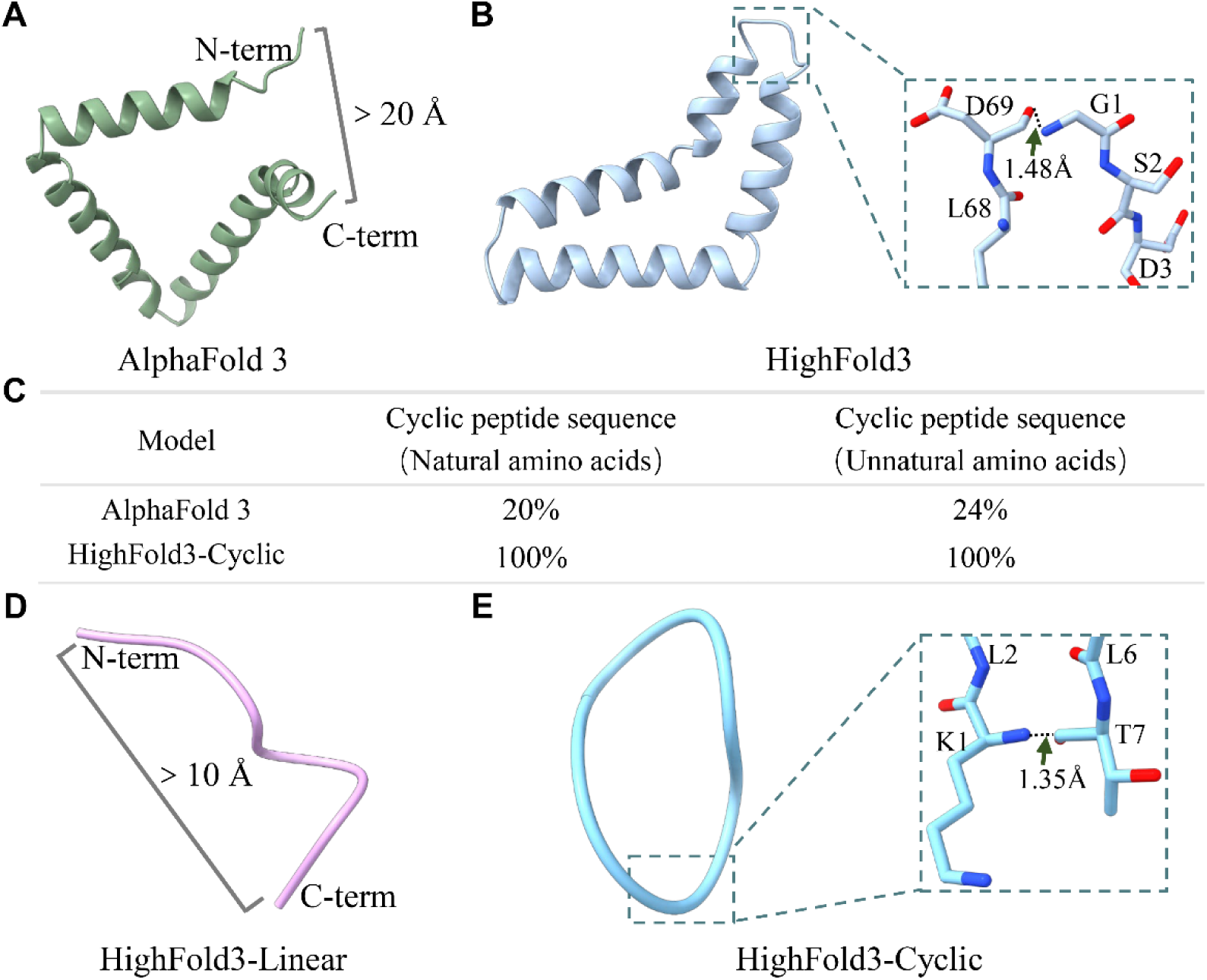
Model Performance in Predicting External Cyclic and Linear Peptide Sequences (A) Overall accuracy of the model on external sequence databases. (B) Structure predicted by AlphaFold3 (PDB ID: 9HVC). (C) Structure predicted by HighFold3 (PDB ID: 9HVC), with a zoomed-in view of the cyclized region. (D) Structure predicted by AlphaFold3 (Sequence: KLARLLT). (E) Structure predicted by HighFold3 (Sequence: KLARLLT), with a zoomed-in view of the cyclized region.

Furthermore, we selected 100 and 50 cyclic peptide sequences from a cyclic peptide database. These sequences, which vary in length and amino acid composition, were obtained from published literature or studies but lack crystal structure data (see Supplementary Files). As shown in Figure 4C, AlphaFold 3 struggles to predict these sequences as closed loops, with success rates of only 20% and 24%, whereas HighFold3-Cyclic module achieves a 100% cyclization success rate. This finding indicates that AlphaFold 3 does not inherently predict cyclic peptides well, whereas HighFold3-Cyclic accurately models cyclic conformations using user-specified cyclization constraints.

The linear or cyclic nature of a peptide does not arise spontaneously in vivo or during in vitro synthesis but is strictly regulated by environmental factors, enzymatic reactions, or artificial catalytic conditions^24^. In nature, cyclic peptides often require non-ribosomal peptide synthetases (NRPS) or post-translational modification enzymes to form stable cyclic structures through head-to-tail linkage or side-chain crosslinking, as seen in the antibiotic vancomycin^25^. Conversely, linear peptides may result from enzymatic cleavage by proteases such as trypsin or pepsin, disrupting an originally cyclic structure, or they may fail to undergo cyclization during synthesis^26^. This dynamic process underscores a key principle: the final conformation (linear or cyclic) of an amino acid sequence is not solely dictated by its sequence but is highly influenced by specific biological or chemical conditions. This insight forms the foundation of HighFold3, which integrates user-specified constraints to accurately distinguish and predict linear and cyclic peptide conformations.

To assess HighFold3’s applicability in practical research, we selected well-documented linear and cyclic peptide sequences from the literature. Williams et al. designed the linear peptide KLARLLT and its cyclic derivative Cyclo(KLARLLT), confirming differences in EGFR binding affinity and serum stability through experiments and constructing their 3D structures via traditional molecular modeling^27^. In this study, we applied HighFold3-Linear and HighFold3-Cyclic to predict their respective structures. As shown in Figure 4D and Figure 4E, HighFold3-Linear predicts KLARLLT as a flexible linear conformation with an N-to-C terminus distance of approximately 10 Å, while HighFold3-Cyclic generates a closed-loop Cyclo(KLARLLT) with an amide bond length of 1.35 Å. To further explore the dynamic interactions between the cyclized peptide and its target, we plan to conduct MD simulations.

These results indicate that AlphaFold 3 has limitations in generalizing beyond its training data, particularly in predicting unseen cyclic peptide structures, whereas HighFold3 offers a more flexible and efficient predictive approach.

## 3. Conclusion

In this study, HighFold3 demonstrated significant advantages in predicting the structures of cyclic and linear peptides containing unAAs. Compared to existing models, HighFold3 achieved higher accuracy in predicting the conformations of complex cyclic peptides, particularly excelling in atomic-level modeling of backbone and side-chain structures. This advancement provides a powerful tool for the rational design of peptide-based drugs, especially for cyclic peptides with enhanced stability and specificity.

The success of HighFold3 is attributed not only to its innovative cyclization constraint mechanism but also to its strong generalization capability for unseen sequences, offering a new tool for studying conformation-dependent peptide functions. However, its prediction accuracy still has room for improvement in certain extreme sequences or complex systems. Compared to traditional experimental methods such as X-ray crystallography and NMR, HighFold3 offers clear advantages in computational efficiency and cost-effectiveness, but its ability to capture dynamic conformations requires further validation. In the future, integrating HighFold3 with MD simulations or multi-scale modeling may further reveal the dynamic behavior of peptides in physiological environments, expanding its applications in biomedicine and drug discovery.

